# In vivo CRISPR screening identifies RNF20/40 as epigenetic regulators of cardiomyocyte maturation

**DOI:** 10.1101/808402

**Authors:** Nathan J. VanDusen, Julianna Y. Lee, Weiliang Gu, Isha Sethi, Yanjiang Zheng, Justin S. King, Ping-Zhu Zhou, Shengbao Suo, Yuxuan Guo, Qing Ma, Guo-Cheng Yuan, William T. Pu

## Abstract

Between birth and adulthood cardiomyocytes (CMs) undergo dramatic changes in size, ultrastructure, metabolism, and gene expression, in a process collectively referred to as CM maturation. The transcriptional network that coordinates CM maturation is poorly understood, creating a bottleneck for cardiac regenerative medicine. Forward genetic screens are a powerful, unbiased method to gain novel insights into transcriptional networks, yet this approach has rarely been used *in vivo* in mammals because of high resource demands. Here we utilized somatic mutagenesis to perform the first reported *in vivo* CRISPR genetic screen within a mammalian heart. We discovered and validated several novel transcriptional regulators of CM maturation. Among them were RNF20 and RNF40, which form a complex that monoubiquitinates H2B on lysine 120. Mechanistic studies indicated that this epigenetic mark controls dynamic changes in gene expression required for CM maturation. These insights into CM maturation will inform efforts in cardiac regenerative medicine. More broadly, our approach will enable unbiased forward genetics across mammalian organ systems.

## INTRODUCTION

At birth, mammalian cardiomyocytes (CMs) undergo maturation, a dramatic and coordinated set of structural, metabolic, and gene expression changes that enable them to sustain billions of cycles of forceful contraction during postnatal life^1,2^. Fetal CMs are primarily glycolytic, mitotic, and contract against low resistance, whereas adult CMs rely on oxidative phosphorylation, are post-mitotic, and support heart growth by increasing in size. Sarcomeric and ultrastructural adaptations, such as plasma membrane invaginations known as T-tubules, facilitate synchronized and forceful CM contraction against high resistance. Unfortunately, CMs induced from stem cells or other non-myocyte sources resemble fetal CMs and lack the hallmark features of mature, adult CMs^3,4^. This “maturation bottleneck” remains a major barrier to using stem cell-derived CMs for disease modeling or therapeutic cardiac regeneration.

The regulatory mechanisms that govern the diverse facets of CM maturation are poorly understood, in large part due to the lack of a suitable *in vitro* model and challenges associated with *in vivo* approaches. Although mosaic gene manipulation strategies have allowed more precise interpretation of *in vivo* experiments with respect to the regulation of maturation^5–7^, the low throughput of standard *in vivo* approaches remains a major barrier. To overcome this obstacle, we sought to perform an *in vivo* forward genetic screen in mice. The resource intensity of traditional forward genetics has precluded their widespread use in mammals, but Cas9 mutagenesis directed by a library of guide RNAs (gRNAs) makes introduction and recovery of gene mutations highly efficient^8–10^, and can be expeditiously deployed mammals *in vivo^5,11,12^*. This capability has been used for forward genetic screens in cultured cells^10,13,14^, but its ability to interrogate endogenous biological processes in mammals *in vivo* has yet to be fully realized.

## RESULTS

### CRISPR loss-of-function screen identifies essential regulators of CM maturation

We developed an *in vivo* forward genetic screen to discover factors that regulate murine CM maturation (Fig. 1a). We employed CRISPR/Cas9 AAV9 (CASAAV) based somatic mutagenesis^5^ and a gRNA library targeting murine transcriptional regulators to create thousands of distinct mutations within different CMs of a single mammalian heart. We screened these mutant CMs with a flow cytometry-based single cell assay of CM maturation. Sequencing of gRNAs from immature CMs compared to the input library identified gRNAs enriched or depleted in immature CMs, i.e. gRNAs that target genes that cell autonomously promote or antagonize CM maturation, respectively.

**Fig. 1:**
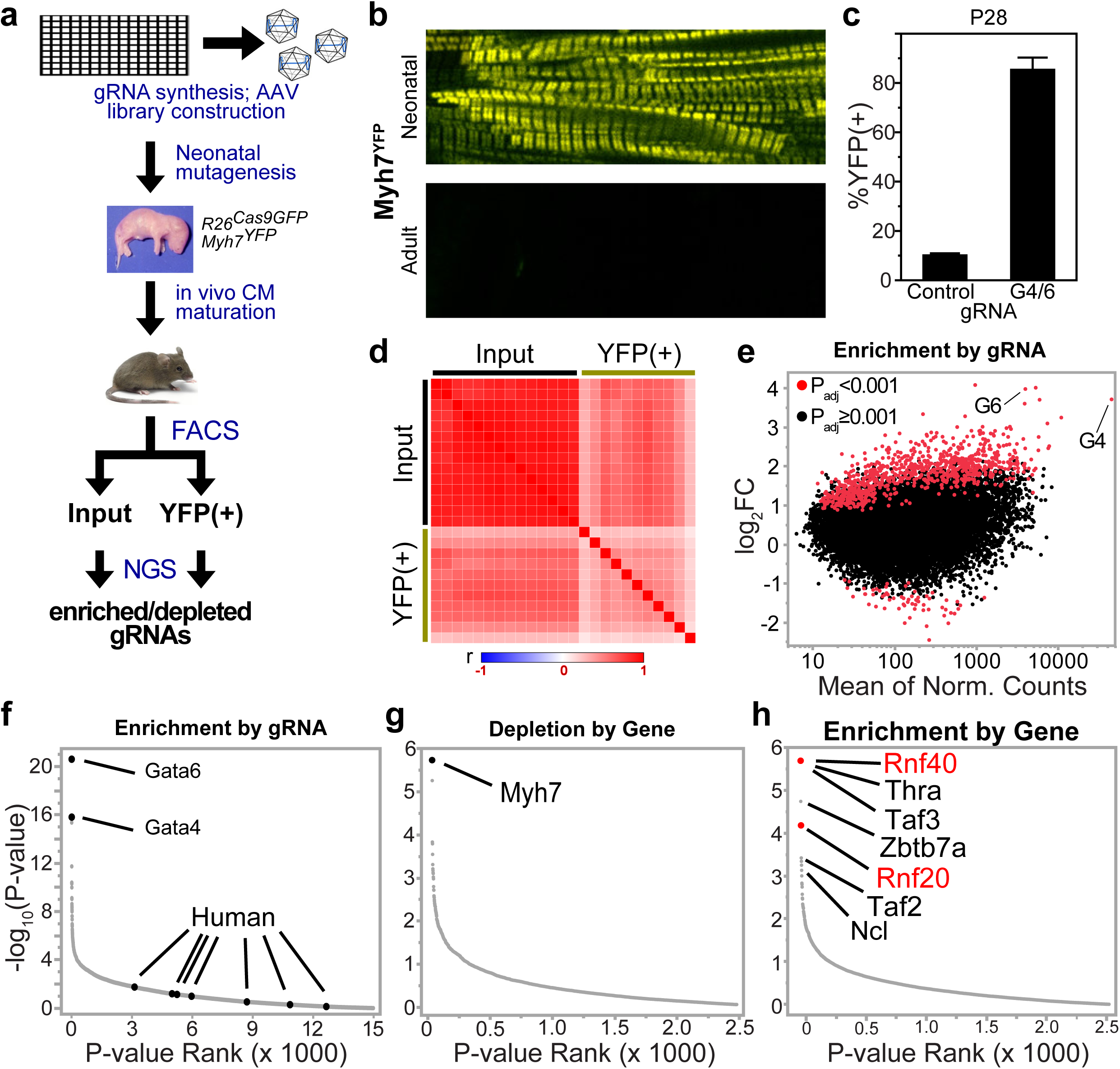
High through-put in vivo CRISPR-Cas9 pooled screen for regulators of cardiomyocyte maturation. **a,** Screen overview. **b,** Myh7^YFP^ expression is restricted to neonatal stage. **c,** Neonatal mosaic double knockout of GATA4 and GATA6 by CASAAV results in persistent MYH7^YFP^ expression. **d,** Sample clustering by r correlation. **e,** DESeq2 differential expression analysis of individual gRNAs. GATA4 and GATA6 positive control gRNAs are indicated. **f,** Enrichment of positive and negative control gRNAs within YFP^+^CMs. **g,** Gene depletion calculated by MAGeCK. **h,** Gene enrichment calculated by MAGeCK

To separate individual CMs with an immature or mature phenotype, we took advantage of developmentally regulated sarcomere gene isoform switching, a hallmark of CM maturation^1,2^. In mouse, a prominent switch is from myosin heavy chain 7 (*Myh7*) expression in the fetal and neonatal periods to *Myh6* in mature CMs. In *Myh7 ^YFP^* mice^15^, YFP is fused to endogenous MYH7, such that YFP fluorescence is controlled by endogenous *Myh7* regulatory elements^15^. We hypothesized that *Myh7^YFP^* could be used as a single cell readout of CM maturation state. Examination of *Myh7^YFP^* myocardium from neonatal and adult mice verified that neonatal CMs exhibited strong YFP fluorescence, whereas YFP was nearly completely silenced in adult CMs (Fig. 1b). Since mutation of both *Gata4* and *Gata6* impair CM maturation (Suppl. Fig. 1 and ref. ^16^), we next tested the effect of *Gata4/Gata6* mutation on *Myh7^YFP^* expression. We found that CASAAV-Gata4-Gata6, which expresses CM specific Cre and gRNAs targeting both Gata4 and Gata6 (Suppl. Fig. 1a), markedly increased the fraction of transduced CMs that expressed YFP from 11% to 86% (Fig. 1c). These data show that *Gata4/6* mutation inhibits normal CM maturation, including maturational *Myh7* silencing.

Because the coordination of CM maturation suggested transcriptional regulation, we focused our screen on transcriptional regulators. We developed a pooled CASAAV library comprised of AAV expressing CM specific Cre and a gRNA designed to target a candidate gene. The candidate gene list contained 1894 transcription factors and epigenetic modifiers. Given that GATA4/6 regulate multiple aspects of CM maturation (Suppl. Fig. 1), we also included 259 genes differentially expressed in P6 Gata4/6 high dose double KO CMs, and 291 genes with strong adjacent G4 binding at P0 (Suppl. Table. 1). In total, 2444 genes were selected for targeting. We used a computational pipeline designed to optimize gRNA on-target activity and yield of frameshift mutations^17,18^ to design six guides for each gene. 7 human gRNAs that do not target the mouse genome were also included as negative controls. These 14671 gRNAs were synthesized as an oligonucleotide pool, cloned into the CM specific CASAAV vector, and packaged into AAV9 (see Methods). To introduce a positive control, a small amount of the CASAAV-Gata4-Gata6 vector was spiked into the AAV pool.

We subcutaneously injected the AAV library into 45 newborn *R26^Cas9-GFP/+^;Myh7^YFP/+^* pups at a dose sufficient to transduce approximately 50% of the myocardium. At four weeks of age mice were sacrificed and CMs isolated. 15% of the isolated CMs from each heart were set aside as an unsorted input sample, while YFP^+^CMs were sorted from the remaining 85% via flow cytometry (Suppl. Fig. 2a). CMs from three hearts were pooled for sorting, resulting in 15 input and 15 YFP^+^ samples. Following RNA isolation, gRNAs were specifically reverse transcribed, converted to barcoded amplicon libraries, and sequenced to an average depth of 4.8M reads. 11 YFP+ and 14 input samples passed quality control (Suppl. Fig. 2b-f), and these samples separated into distinct clusters (Fig. 1d). 834 gRNAs showed significant enrichment within YFP^+^ samples (adj. *P* < 0.001), with the *Gata4* and *Gata6* positive control gRNAs being among the most enriched (Fig. 1e). The seven human-targeting negative control gRNAs did not show enrichment (Fig. 1f). We used MaGeCK^19^ to consolidate the six gRNA enrichment scores per gene into a single score. gRNAs targeting 123 genes were significantly enriched within YFP^+^CMs, while gRNAs targeting 148 genes were depleted (*P* < 0.05). The top ranked depleted gene was *Myh7* (Fig. 1g), which was expected given that YFP fused to *Myh7* was the screen readout. Among the enriched genes were thyroid hormone receptor alpha (*Thra*) and nucleolin (*Ncl*), which are established regulators of maturation ^20,21^, as well as many novel candidates (Fig. 1h; Suppl. Table 2), including both *Rnf20* and *Rnf40*, which encode components of an epigenetic complex that monoubiquitinates histone 2B^22,23^.

**Fig. 2:**
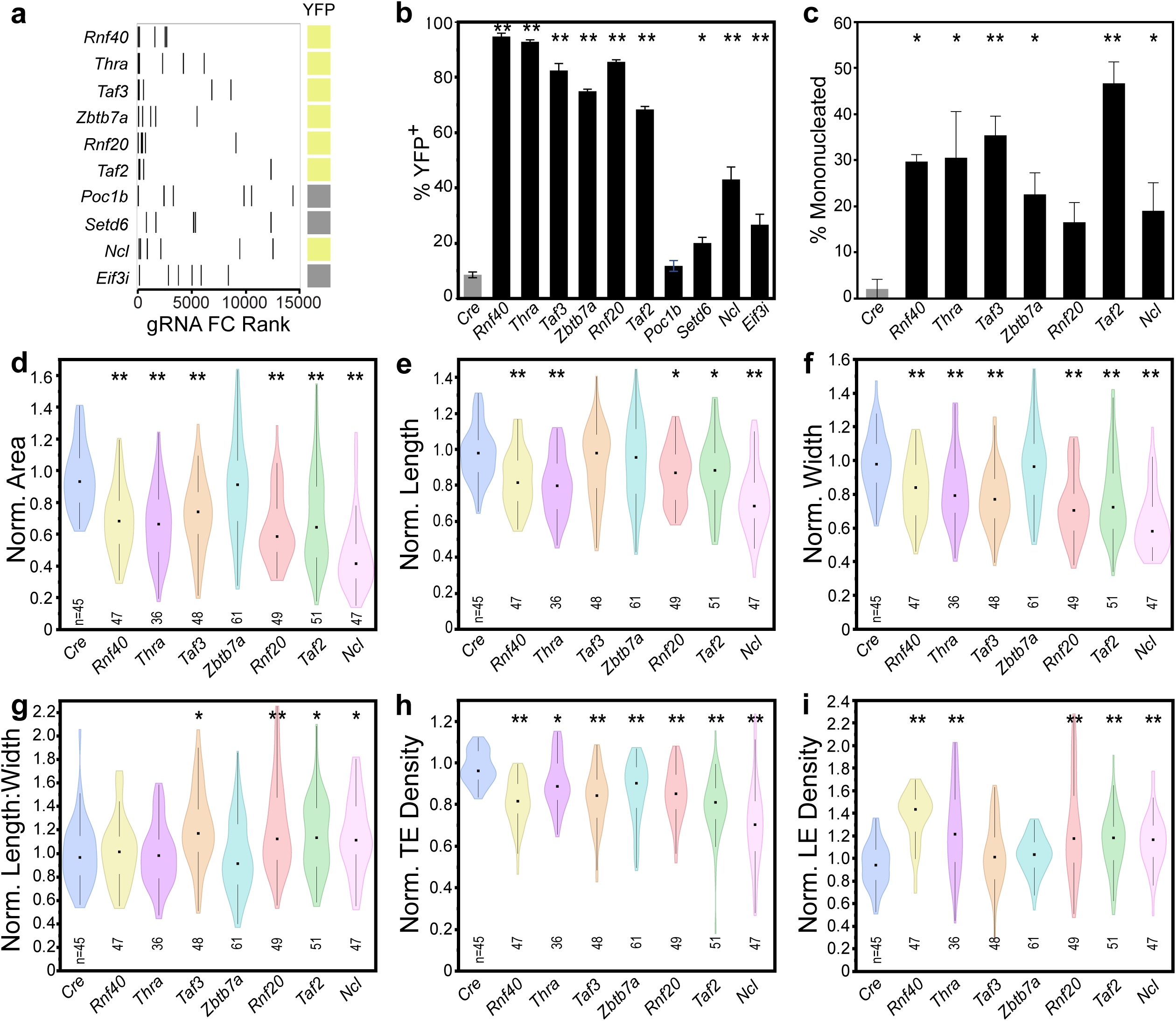
Validation of top candidates. **a,** Top ten candidates ranked high to low by MAGeCK enrichment score. The six guides for each candidate are ranked by individual enrichment, where higher enriched gRNAs have a smaller rank. Candidates marked in yellow induced robust persistent activation of Myh7^YFP^ when individually targeted by the most highly enriched gRNA. **b,** Quantification of Myh7^YFP^ activation within transduced (GFP+) CMs. n = 3. **c,** Quantification of mononucleation among GFP+ control CMs or YFP+ candidate depleted CMs. n = 3. **d-g,** Normalized length, width, length to width ratio, and area for control and candidate depleted CMs. Measurements from YFP+ CMs are normalized to YFP-cells from the same heart. **h-i,** Normalized T-tubule transverse element density and longitudinal element density for control and candidate depleted CMs as measured by AutoTT software. Bar plots show mean ± SD. Student’s t test: \**P*<0.05, \*\**P*<0.001

To validate the top 10 most enriched candidates (Fig. 2a), the most highly enriched gRNA for each factor was cloned into a CASAAV vector and used to individually deplete each factor at birth. To focus on cell autonomous effects and avoid secondary effects related to organ dysfunction^5–7^, we used an AAV dose that transduced a small fraction of CMs. Among mosaic depleted cells, we observed robust upregulation of *Myh7^YFP^* for seven of the ten candidates: *Rnf40, Rnf20, Taf3, Taf2, Thra, Zbtb7a* and *Ncl* (Fig. 2a,b; Suppl. Fig. 3a). CASAAV vectors targeting the remaining three candidates, *Poc1b, Setd6*, and *Eif3i*, caused little YFP activation (Fig. 2a,b, Suppl. Fig. 3a). Consequently, we focused subsequent analyses on the seven target genes that resulted in robust upregulation of *Myh7^YFP^*.

**Fig. 3:**
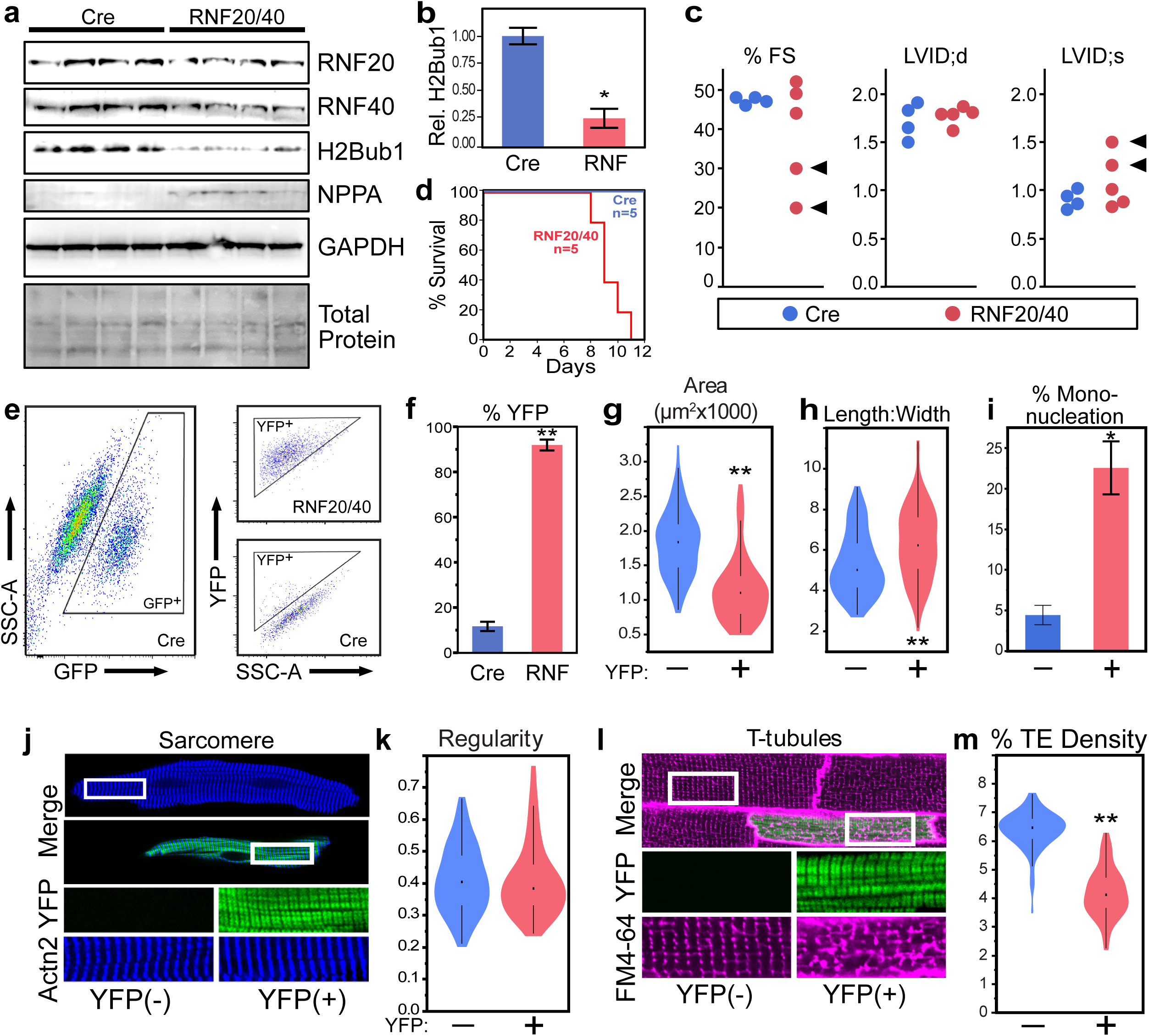
Depletion of RNF20/40. **a,** Western blot of P7 cardiac apex from pups transduced with a high dose of control or RNF20/40 gRNAs at birth. **b,** Quantitation of H2Bub1 depletion. **c,** Echocardiography of P7 mouse pups injected with a high dose of RNF20/40 targeting gRNAs at birth. Arrows indicate animals with cardiac dysfunction. **d,** Survival curve of mice receiving a high dose injection of RNF20/40 CASAAV. **e,** Representative gating of transduced CMs. Representative gating for quantitation of persistent Myh7^YFP^ expression in mosaic transduced CMs. **f,** Quantitation of sustained Myh7^YFP^ expression in control or RNF20/40 mosaic double depletion CMs. Cre n=3, RNF20/40 n=4. **g,** Area of control or YFP+ adult CMs from mice transduced at birth with mosaic dose of RNF20/40 targeting gRNAs. **h,** Length to width ratio. **i,** Percent of cells mononucleated. **j,** Representative images of sarcomere organization as visualized by sarcomeric alpha actinin staining. **k,** Quantitation of sarcomere organization. **l,** Representative images of t-tubule organization as visualized by plasma membrane binding dye FM4-64. **m,** Quantitation of t-tubule transverse element density. Bar plots show mean ± SD. Student’s t test: \**P*<0.05, \*\**P*<0.001.

We analyzed the effect of individual CASAAV vectors on CM nucleation, size, and T-tubulation, additional hallmarks of maturation. Immature murine CMs are mononuclear and become predominantly binuclear during maturation^24^, and this polyploidization has been implicated in the reduced regenerative potential of mature CMs^25–27^. CMs markedly increase in cell size between birth and adulthood^28^, as cellular hypertrophy accounts for the postnatal increase in heart size required to meet intensified postnatal demands. T-tubules, highly organized invaginations of the cell membrane, are necessary for fast and synchronized excitation-contraction coupling in large mature CMs. T-tubules develop in mice by the end of the second postnatal week^29^. The seven CASAAV vectors or AAV-Cre lacking gRNA (control) were delivered individually to newborn mice at a mosaic dose. At one month of age, CMs were dissociated, fixed, and stained to visualize T-tubules (CAV3) and nuclei (DAPI). Results were compared between YFP^+^ CMs and control CMs, marked by GFP. Five of the seven CASAAV vectors (*Rnf40, Thra, Taf3, Taf2*, and *Ncl*) increased mononucleation, decreased cell size, and disrupted T-tubules (Fig. 2c-i; Suppl. Fig. 3a,b). CASAAV-Rnf20 impaired maturational growth and T-tubulation, but the increase in mononucleation did not reach statistical significance (*P*=0.0763). *Zbtb7a* increased mononucleation and impaired T-tubulation, but did not influence maturational hypertrophy. These results validated 7 of the 10 candidates as regulators of multiple facets of CM maturation, and also demonstrate that the overall maturational program can be separated into independently regulated, dissociable sub-programs.

### RNF20/40 depleted CMs display morphological and transcriptional characteristics of immaturity

*Rnf20* and *Rnf40* were two of the most enriched genes in the screen, and their depletion broadly impaired CM maturation. These genes are E3 ubiquitin ligases that together form a complex that monoubiquitinates histone 2B at lysine 120 (H2Bub1)^30,31^. Human genetic studies have implicated *de novo Rnf20*/*Rnf40* mutations in congenital heart disease^32,33^. The postnatal cardiac functions of these genes have not been studied. For these reasons, we investigated the mechanisms by which RNF20/40 regulate CM maturation. We created a single CASAAV vector (CASAAV-RNF20/40) containing gRNAs that target both factors and validated that it depleted RNF20/40 and H2Bub1 (Fig. 3a,b). When given at a dose that transduced most CMs, CASAAV-RNF20/40 caused cardiac dysfunction and death (Fig. 3c,d). At a dose that transduced a small fraction of CMs (Fig. 3e), RNF20/40 depletion cell autonomously impaired maturational silencing of Myh7^YFP^ (Fig. 3e,f), maturational growth (Fig. 3g,h), and maturational multi-nucleation (Fig. 3i). While sarcomere organization was unaffected (Fig. 3j,k), we observed dramatic defects in T-tubule organization (Fig. 3i,m).

We next performed transcriptional profiling to measure the effect of RNF20/40 depletion on gene expression. Newborn *R26^fsCas9-2A-GFP/+^;Myh7^YFP/+^* mouse pups were injected with a mosaic dose of CASAAV-RNF20/40 vector, or a control vector containing Cre without gRNAs. At P28 hearts were dissociated and flow cytometry was used to recover YFP^+^CMs transduced with CASAAV-RNF20/40 or control GFP^+^CMs transduced with Cre. RNA from each group was isolated and transcriptomes analyzed by RNA-seq (Fig. 4a). Principal component analysis showed clear separation of sample groups (Fig. 4b), and differential gene expression analysis revealed approximately 1400 upregulated and 1100 downregulated genes (*P*_*adi*_ < 0.05, Fig. 4c, and Suppl. Table 3). The ratio of fetal (*Tnni1*) to mature (*Tnni3*) troponin I isoforms, a molecular signature of CM maturational state^34^, was increased by 165-fold in RNF20/40 depleted CMs (*P* < 0.001), with *Tnni1* being the most upregulated gene in the dataset. To assess the genome wide impact of RNF20/40 depletion we used RNA-sequencing from neonatal and adult isolated wildtype CMs to construct custom gene sets consisting of the 100 most neonatal specific, and 100 most adult specific genes. We then ranked all genes by their level of differential expression in the RNF20/40 depleted CMs, and used Gene Set Enrichment Analysis (GSEA)^35^ to calculate an enrichment score for each custom gene set within this ranked list. The genes downregulated in RNF20/40 depleted CMs were highly enriched for adult-specific genes (NES = −1.82; Fig. 4d), while the genes upregulated in the depleted CMs were enriched for neonatal-specific genes (NES = 1.50; Fig. 4e). These data confirm that RNF20/40 depleted cells fail to activate the transcriptional network of mature CMs, but instead persistently express genes associated with immaturity.

**Fig. 4:**
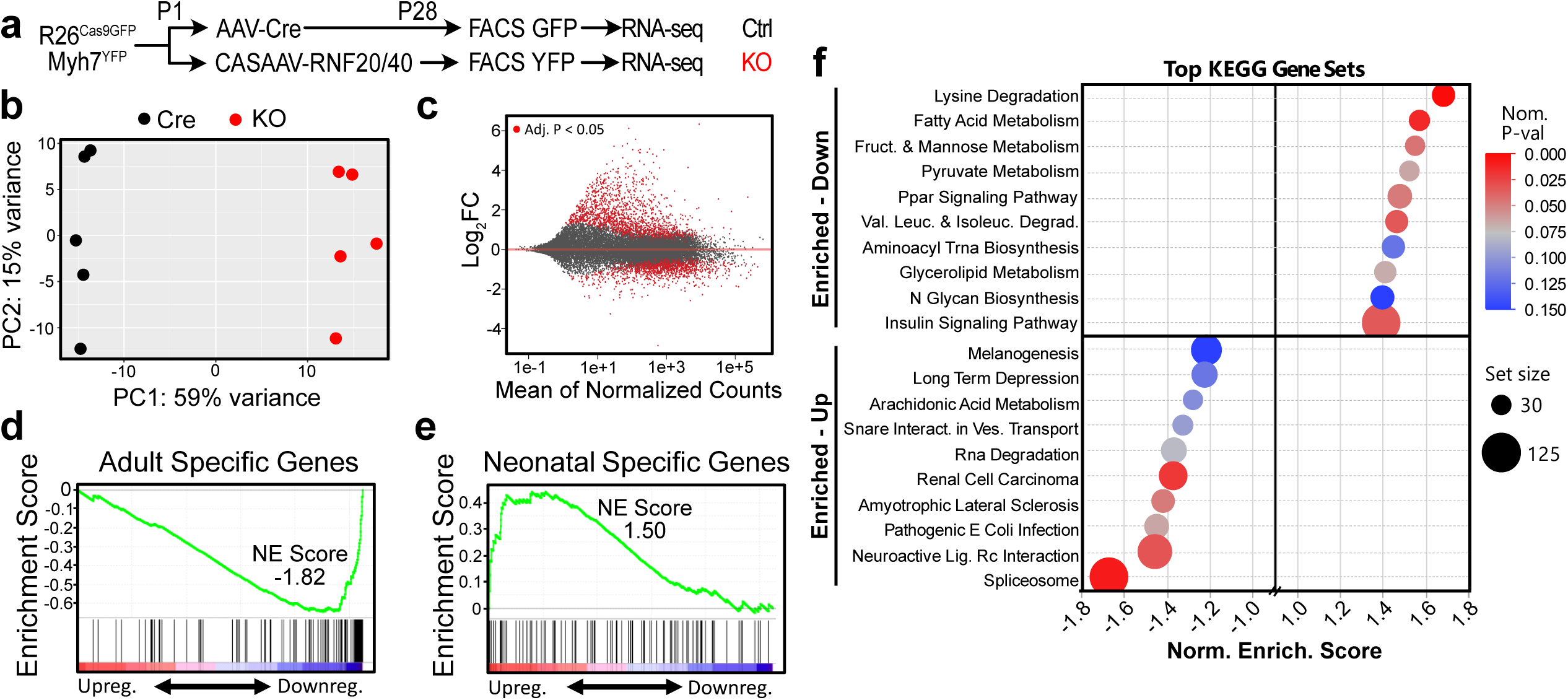
Transcriptional profiling of mosaic RNF20/40 depleted CMs. **a,** Experiment overview. **b,** Principal component analysis of RNA-seq replicates. **c,** Differentially expressed genes. **d,** GSEA with a custom “Adult Specific Genes” gene set. **e,** GSEA with a custom “Neonatal Specific Genes” gene set. **f,** GSEA of gene list ranked by differential expression in RNF20/40 KO CMs.

We next analyzed the RNA-seq data to identify biological processes enriched within the RNF20/40 differentially expressed genes. Many metabolism-related gene sets were enriched within the downregulated genes (Fig. 4f, Suppl. Table 5), including Fatty Acid Metabolism and PPAR Signaling, two pathways strongly associated with CM maturation^36,37^. Upregulated genes were enriched for a greater diversity of biological processes, with spliceosome-associated genes being most enriched (Fig. 4f).

### H2Bub1 deposition directly regulates transcriptional maturation

The RNF20/40 complex is an E3 ubiquitin ligase that monoubiquitinates H2B on lysine 120 (H2Bub1). To determine how H2Bub1 deposition by RNF20/40 promotes normal maturation of CM gene expression, we used chromatin immunoprecipitation followed by next generation sequencing (ChIP-seq) to determine the genomic distribution of H2Bub1 at neonatal (P1) and mature stages (P28) in mouse heart apex tissue. Consistent with prior reports^38^, H2Bub1 predominantly occupied gene bodies, with greater density towards promoters. Thus, we quantified the amount of H2Bub1 deposition at each gene and observed reproducible signal above input at approximately 11,400 genes in the neonatal stage and 11,800 genes in the adult stage. Of these genes, the vast majority (>88%) were shared between timepoints (Fig. 5a), with a small number of unique regions (Fig. 5b, representative genome browser view). At either stage, genes with greater H2Bub1 generally were more highly expressed, although the correlation was poor (R^2^∼0.01;Fig. 5c). However, there was a significant correlation between the change in H2Bub1 and the change in gene expression between stages, consistent with the reported association of H2Bub1 with gene activation (Fig. 5d). Strikingly, genes with greater H2Bub1 were less likely to substantially change expression between stages (Fig. 5e). This stabilizing effect of H2Bub1 was also observed when comparing RNF20/40 depleted CMs to negative controls (Fig. 5f), suggesting that high H2Bub1 induces additional epigenetic mechanisms that limit changes in gene expression.

**Fig. 5:**
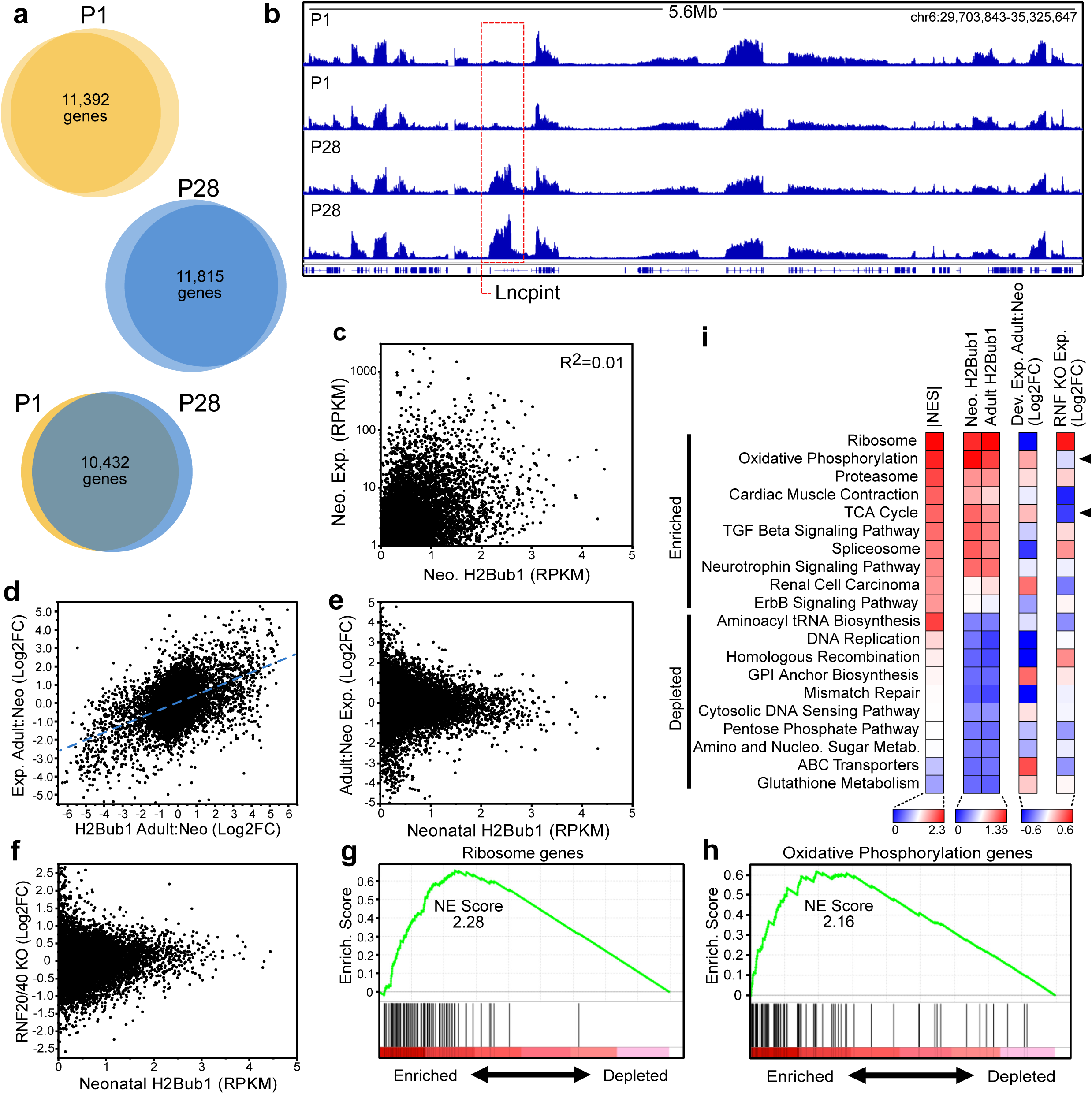
RNF20/40 directly regulate metabolism during CM maturation. **a,** Number of genes marked by H2Bub1 at neonatal and adult timepoints. Two ChIP-seq replicates conducted for each timepoint. **b,** Genome browser view containing one region with gain of H2Bub1 during maturation. **c,** Relationship between gene expression and H2Bub1 deposition at neonatal timepoint. **d,** Relationship between change in expression and change in H2Bub1 deposition during maturation. Dashed line is best fit line. **e,** Relationship between expression change during maturation and level of H2Bub1 deposition at neonatal stage. **f,** Relationship between expression change in RNF20/40 depleted CMs and level of H2Bub1 deposition at neonatal stage. **g,** GSEA showing high deposition of H2Bub1 at ribosomal genes in neonatal stage. **h,** GSEA showing high deposition of H2Bub1 at oxidative phosphorylation genes in neonatal stage. **i,** Top 10 most enriched and most depleted gene sets, with heatmaps of H2Bub1 signal at neonatal and adult stages, gene expression change during maturation, and gene expression change in RNF20/40 depleted CMs. Arrows indicate metabolic gene sets that show high enrichment within H2Bub1 ranked list, activation during CM maturation, and loss of expression in RNF20/40 depleted CMs.

Next we used GSEA to identify biological processes containing genes preferentially marked by H2Bub1. Several functional terms were associated with strong H2Bub1 signal at the neonatal stage (Nom. Pval < 0.001, NES > 1.75), with Ribosome and Oxidative Phosphorylation being the top two (Fig 5g,h). While most genes highly marked by H2Bub1 have a stable expression profile during maturation (Fig. 5e), we observed that genes within several of the identified functional groups are dynamically expressed during maturation, and differentially expressed in RNF20/40 CMs (Fig. 5i). This included Oxidative Phosphorylation and TCA Cycle gene sets, suggesting that H2Bub1 deposition is directly required for activation of the adult metabolic gene profile.

## DISCUSSION

In this study we developed a resource efficient *in vivo* forward genetic screen for transcriptional regulators of CM maturation. Our strategy uses a massively parallel approach in which the screening unit is the individual cell, such that thousands of individual genetic mutations can be screened using a small number of animals. Among the many novel genes that our screen identified as essential for CM maturation were *Rnf20* and *Rnf40*, which interact to form a complex that mono-ubiquitinates histone-2B. Depletion of *Rnf20/40* broadly impaired CM maturation by disrupting H2Bub1 deposition, which prevented normal transcriptional changes essential for maturation. While *Rnf20/40* regulate diverse gene sets, one common theme in our studies was regulation of the metabolic remodeling that takes place during CM maturation. GSEA identified lysine degradation and fatty acid metabolism as the two genesets most downregulated in RNF20/40 depleted CMs, while an independent GSEA based on H2Bub1 deposition levels identified oxidative phosphorylation and TCA cycle gene sets as preferentially marked.

Mutations in pathways that regulate H2Bub1, including *Rnf20* and *Rnf40*, have been reported in humans with congenital heart defects (CHD)^32,33^ and were recently reported to influence cardiac looping via regulation of motile cilia during early embryonic development^32^. Our results extend the function of *Rnf20/40* into the postnatal heart, suggesting that mutations in these genes could both cause a structural heart defect and contribute to cardiac dysfunction after surgical repair. CHD survivors have a greatly increased risk of cardiovascular disease later in life^39^, with outcomes being diverse and difficult to predict. As there is substantial diversity in the range of CHD genes, additional research will be necessary to characterize the role of these factors in CM maturation, to discover if functional declines are due to cell autonomous maturational defects, and to inform innovations in intervention.

## MATERIALS AND METHODS

### Library Design and Construction

A list of 3000+ potential transcription factors and epigenetic regulators was compiled using publicly available data from the Riken Transcription Factor Database^40^ and AnimalTFDB^41^). RNAseq data from isolated P6 wild type CMs was used to determine which factors were expressed in the neonatal mouse heart. Factors with an average FPKM >1 across three replicates were included in the screen (1735 regulators). 159 randomly selected regulators with an FPKM of 0 were included in the library as negative controls. In addition, 259 genes differentially expressed at P6 within GATA4/6 double KO CMs (adjusted p-value < 0.05), and 291 genes with strong adjacent GATA4-fbio binding at P0 were also included (GEO - GSE124008), for a total of 2444 genes targeted. Six gRNAs were selected for each gene, using CRISPR RGEN Tools Cas-Database^18^. The following five rules were applied, in order, to select the “best” six guides per gene (ie. proceed to next rule if less than six guides are available which meet the current rule). 1) gRNAs target constitutive exons, CDS range 5-50%, number of mismatches 0,1,2:1,0,0 meaning only one on-target exact match, zero off target sites with one mismatch, and zero off-target sites with two mismatches, and microhomology-associated out of frame score >60. 2) gRNAs target constitutive exons, CDS range 5-80%, number of mismatches 0,1,2:1,N,N and out of frame score >60. 3) gRNAs target any exon, CDS range 5-80%, number of mismatches 0,1,2:1,N,N and out of frame score >60. 4) gRNAs target any exon, CDS range 5-80%, number of mismatches 0,1,2:N,N,N and out of frame score >60. 5) gRNAs target any exon, any CDS range, number of mismatches 0,1,2:N,N,N and any out of frame score. Approximately 63% of guides in the library met rule one criteria. A 5’ G was added to each gRNA to ensure optimal expression from the U6 promoter, and gRNAs were flanked by SapI restriction sites. The 14,664 guide library was synthesized by Agilent as a single 80bp SUREprint oligo pool (Supplemental Table 6). The library was resuspended in 50ul of TE (200nM) and diluted to 33nM. 1ul of 33nM library was amplified for 10 cycles via a standard NEB Phusion PCR program, and 80bp library amplification primers (Supplementary Table 7), to produce a 200bp amplicon. Eight reactions were pooled, cleaned up via a Zymo Research DNA Clean and Concentrator Kit (Zymo, D4014), and 5ugs were digested with SapI for 3 hours. The 21bp gRNA library was purified via Invitrogen Size Select 2% gel, and seamlessly ligated into our previously described CASAAV vector (Addgene 132551)^5,42^. Ligation product was purified via Zymo DNA column, and electroporated into Agilent SURE Electrocompetent cells with a Bio-Rad Gene Pulser Xcell Electroporation System. 40ng of purified ligation product in 2ul volume was electroporated into 40ul of cells. Four electroporations were conducted for a total of 160ng of ligation product. Electroporations were conducted at 1700V, 200 ohms resistance, 25μF capacitance, and 1mm cuvette gap. 900ul of SOC media was added immediately after electroporation, and bacteria were incubated at 37°C for 1hr. Bacteria were plated on LB agar containing ampicillin and allowed to grow for 18hrs. Approximately 300,000 colonies (20x library coverage) were scraped into SOC media, cultured for additional 1.5hrs, and plasmid DNA harvested (Invitrogen Purelink HiPure Maxiprep, 210017), yielding 240ug of DNA. This library pool was packaged into AAV9 via PEI transfection of ten 15cm plates of HEK293T cells, and titered by qPCR targeting the TnT promoter, as previously described^5,42^.

### Injections, Sample Collection, and Flow Cytometry

One day old R26^fsCas9-2A-GFP/+^;Myh7^YFP/+^ pups were subcutaneously injected with 50ul of gRNA library virus at a concentration of 2×10^11^vp/ml, spiked with a single CASAAV virus targeting both GATA4 and GATA6 at a final concentration of 1×10^9^vp/ml. At four weeks of age, mice were sacrificed and single cell CM dissociations prepared by collagenase perfusion, as previously described^5,42^. Atria were removed and discarded after perfusion, prior to dissociation. 15% of the isolated CMs (∼200,000) were set aside as an unsorted input sample, while YFP^+^ CMs were sorted from the remaining 85% via FACS. Sorting was performed at the DANA Farber Cancer Center flow cytometry core on an Aria II cell sorter with 100um nozzle, and with 510/21 bandpass filter for GFP, 550/30 bandpass filter for YFP, and 525 longpass dichroic filter to split GFP and YFP signals. CMs were sorted into Trizol (Life Technologies, 15596026), and along with input samples, RNA was extracted via standard phase separation. CMs from three hearts were sorted into each RNA collection tube (45 hearts, 15 pooled samples). Trizol aqueous phase was transferred to a Zymo RNA Clean and Concentrator spin column (Zymo, R1015) and treated with DNase (Qiagen, 79254) for 20 minutes, followed by cleanup, and elution in 15ul of H2O for YFP samples and 120ul of H2O for Input samples. After isolation, Input RNA samples were pooled in groups of three to match YFP samples.

### NGS Library Preparation

Reverse transcription of gRNAs and adapter addition was achieved by using a custom protocol with the Clontech SMART-Seq v4 Ultra Low Input RNA sequencing kit (634894). Approximately 30ng of RNA from YFP^+^ CMs or 500ng of RNA from Input CMs was reverse transcribed as directed in the manufacture protocol except that SMART-Seq CDS Primer IIA was replaced with a gRNA scaffold specific primer (Supplementary Table 7). This reverse transcription step also utilizes a template switching oligo to add an adapter of known sequence to the variable 5’ end of the gRNA^43^. cDNA was then amplified in two sequential rounds of PCR to add NGS sequencing adapters. In the first round of amplification the full length forward read adapter was added to the 5’ end of the gRNA and a half adapter was added to the 3’ end (Supplementary Table 7). NEB Phusion (M0530L) was used for 5 cycles of amplification, according to standard manufacturer protocol. First round PCR product was purified via Zymo DNA Clean and Concentrator column, eluted in 25ul of H2O, and 5ul used as input in a 20 cycle second round of amplification, which completed the reverse adapter and added a sample specific Illumina TruSeq multiplexing index. The resulting 220bp amplicon was purified via Invitrogen SizeSelect 2% gel. The concentration of each sample was assessed using the KAPA Library Quantitation Kit (KR0405), allowing samples to be evenly pooled and submitted for single end 75bp sequencing on a NextSeq500. An average sequencing depth of 4.8 million reads per sample was achieved.

### Screen NGS Analysis

gRNA NGS libraries were trimmed to remove adapters and the gRNA scaffold, leaving only the 20bp variable region. Bowtie2 was used to align trimmed sequence files to mm10. Counts for each gRNA were acquired by quantifying sequence coverage of genomic regions corresponding to the gRNA library via Bedtools Coverage. Differential expression of individual gRNAs in YFP^+^ versus Input samples was calculated by using gRNA counts as input for DESeq2^44^. Differential gene representation in YFP^+^ versus Input samples was calculated using median normalized gRNA counts as input for the MAGeCK software package^19^, which consolidates scores for multiple individual gRNAs targeting the same gene into a single gene level enrichment score. Five samples were removed prior to MAGeCK analysis due to insufficient enrichment of control gRNAs, improper clustering, or poor library coverage (Suppl. Fig. 2).

### In Situ T-tubule Imaging

CASAAV virus targeting *Rnf20* and *Rnf40* for double depletion was injected subcutaneously at P1 into R26^Cas9/+^;Myh7^YFP/+^ pups. Animals were sacrificed at P28 and hearts were perfused with the T-tubule binding dye FM 4-64 (Thermo, T3166) at 5uM for 20 minutes, followed by imaging of on an Olympus FV3000R confocal microscope, as previously described^5^. Organization and abundance of transverse and longitudinal T-tubule elements was quantified using AutoTT software ^45^. In total 79 YFP^-^ and 69 YFP^+^ CMs, originating from three mice, were quantified.

### Immunostaining

Immunostaining was conducted as previously described^42^. Briefly, freshly isolated adult CMs were cultured on laminin coated (2ug/cm^2^, Life Technologies, 23017015) 12mm glass coverslips in a 24-well dish with DMEM plus 5% FBS and 10uM Blebbistatin (EMD Millipore, 203390) at 37°C. After allowing 30 minutes for cells to adhere to the laminin, CMs were fixed with 4% PFA for 10 minutes at room temperature, followed by permeabilization with PBST (PBS + 0.1% Triton) for 10 minutes at room temperature. Cells were then blocked with 4% BSA/PBS for 1hr, and incubated with primary antibody (Supplementary Table 8) diluted 1:500 overnight at 4°C. The next day CMs were briefly rinsed three times with 4% BSA/PBS and incubated with fluorescently conjugated secondary antibodies for 1 hour at 4°C. After rinsing three times with 4% BSA/PBS, coverslips were mounted on slides using Diamond Antifade mountant (Thermo, P36965), and imaged on an Olympus FV3000R confocal microscope.

### Western Blotting

P7 CASAAV RNF20/40 depleted hearts were homogenized in 1ml of RIPA buffer and agitated at 4°C for 30 minutes. Lysates were spun at 10,000g for 10 minutes at 4°C, and the supernatant transferred to a new tube. 40ug of protein was boiled in 2x SDS loading buffer for 5 minutes, and loaded onto an Invitrogen Bolt 4-12% gradient precast mini-gel (NW04120BOX) and run at 165 volts for 45 minutes. Protein was transferred to a pre-equilibrated PVDF membrane in Boston BioProducts transfer buffer (BP-190) via Biorad Trans-Blot SD Semi-Dry Transfer Cell, at 20 volts for 40 minutes. Blots were cut and blocked in 2% milk/TBST for 1 hour at 4°C. Blots were then incubated overnight at 4°C with primary antibodies for H2Bub1, Nppa, and Gapdh, in block solution. See Supplementary Table 8 for antibody product information and dilutions. The next day blots were rinsed in TBST and incubated with HRP conjugated donkey anti-rabbit secondary antibody in blocking solution for 1 hour at room temperature. Blots were then rinsed in TBST, incubated in Millipore Immobilon Western Chemiluminescent HRP Substrate (WBKLS0500) for 1 minute, and imaged on a GE Healthcare ImageQuant LAS4000.

### Echocardiography

Echocardiography was performed on a VisualSonics Vevo 2100 machine with the Vevostrain software. Animals were awake during this procedure and held in a standard handgrip. The echocardiographer was blinded to genotype and treatment.

### RNA-sequencing

One day old R26^fsCas9-2A-GFP/+^;Myh7^YFP/+^ pups were subcutaneously injected with 50ul of CASAAV, containing either the most enriched Rnf20 targeting gRNA and the most enriched Rnf40 targeting gRNA in a single vector (CASAAV-Rnf20-Rnf40), or a control vector containing Cre but no gRNAs. Vectors were injected at a concentration of 1×10^11^vp/ml, which was sufficient for ∼20% transduction. At four weeks of age mice were sacrificed and single cell CM dissociations prepared by collagenase perfusion. After perfusion, ventricular apexes were dissected out and dissociated. YFP^+^RNF20/40 depleted CMs, or control GFP^+^CMs (without regard for YFP), were sorted into Trizol and RNA was extracted via standard phase separation. Approximately 100,000 CMs were collected from each of five RNF20/40 depleted and five control mice. Trizol aqueous phase was transferred to a Zymo RNA Clean and Concentrator spin column and treated with DNase prior to elution in 20ul H2O, as described above. Reverse transcription of 40ng of RNA for each sample, and library amplification, was achieved using the SMARTseq v4 Ultra Low Input RNA-seq kit (Clontech 634889). The standard manufacturer protocol was followed, with 8 cycles of amplification. To prepare amplified libraries for sequencing, 300 pg of each sample was enzymatically fragmented and indexed using a Nextera XT DNA Library Preparation Kit (Illumina FC-131-1024), and Index Kit (Illumina FC-131-1001), according to the standard manufacturer protocol. Single end 75 bp sequencing of pooled libraries was performed on a NextSeq500. After trimming the first 15 bp from sample reads to remove adapter sequences, reads were aligned to a fasta file of the mm10 transcriptome (ftp://ftp.ensembl.org/pub/release-93/fasta/mus_musculus/cdna/) using Kallisto^46^. An average sequencing depth of ∼32M pseudoaligned reads per sample was achieved. Kallisto counts for individual transcripts were consolidated into gene level counts using TxImport^47^, and analyzed for differential expression between control and RNF20/40 depleted sample groups using DESeq2 (Suppl. Table 3)^44^. Gene ontology analysis of differentially expressed genes was conducted with Gene Set Enrichment Analysis^35^, with 1000 permutations and reported gene sets being limited to those containing at least 30 genes (Supplementary Table 5). For construction of custom gene sets RNA-sequencing data from P0 and P7 isolated mouse CMs (GEO - GSE124008) was averaged to get a “neonatal” RPKM expression value, and data from 4 week and 6 weeks of age averaged to get an “adult” value. Genes were then ranked based on the adult to neonatal ratio, and the highest 100 genes selected for an “adult specific gene set” and the lowest 100 for a “neonatal specific gene set” (Suppl. Table 4).

### ChIP-sequencing

Each final ChIP-seq sample consisted of 20 apexes from bisected P1 hearts or 4 apexes from P28 hearts. Each sample consisted of male and female tissue in equal proportions. Tissue, either 2 adult apexes or one litter of P1 apexes, was harvested into ice cold PBS, transferred into 1.2ml of 1% formaldehyde (Sigma F8775), and homogenized with a T10 Ultra Turrax homogenizer at setting 6 for 30 seconds. Homogenate was crosslinked at room temperature on a rotator for 25 minutes. Formaldehyde was quenched by adding glycine to a final concentration of 500mM, and incubating for 5 minutes at room temperature with rotation. Homogenate was then centrifuged at 3000g for 3 minutes at 4°C, supernatant discarded, and tissue resuspended in 1 mL of cold PBS. This PBS wash was repeated for a total of three times, with the tissue pellet being resuspended in hypotonic buffer (20mM HEPES pH 7.5, 10mM KCl, 1mM EDTA, 0.1mM activated Na3VO4, 0.5% NP-40, 10% glycerol, 1mM DTT and 1:1000 Roche cOmplete protease inhibitor) after the third wash. Resuspended tissue was transferred into a glass douncer and dounced with pestle “B” (20x for neonatal tissue, 200x for adult). Lysed cells were transferred into siliconized Eppendorf tubes and incubated on ice for 15 minutes. Lysates were centrifuged at 13,000g for 2 minutes at 4°C, and supernatant discarded. Pellets were stored in liquid nitrogen until multiple samples were ready for sonication. Prior to sonication, pellets were thawed, resuspended in 1ml of hypotonic buffer per 20 neonatal apexes or 2 adult apexes, and incubated on ice for 5 minutes. Lysate was transferred into glass douncer, and again dounced with pestle “B” (20x for neonatal tissue, 200x for adult). Lysate was centrifuged at 13,000g for 2 minutes at 4°C, and the supernatant was discarded. The pellet was then resuspended in 500ul of ChIP Dilution Buffer (20mM Tris-Cl pH 8.0, 2mM EDTA, 150mM NaCl, 1% Triton X-100) supplemented with SDS to 1%, and 1:50 protease inhibitor. Samples were then equally split into two 0.65mL tubes (BrandTech 781310 or Corning 3208) for sonication. Samples were sonicated using a Qsonica 800R, at 65% amplitude, 10 seconds on, 30 seconds off, for 25 minutes of on time (100 minutes total). After sonication the half samples were recombined in 1.5ml siliconized tubes and centrifuged at 18,500g for 5 minutes at 4°C. The chromatin/supernatant was transferred to a new tube, with adult samples being combined so that each tube represents 4 apexes. A 20ul aliquot was removed as an input sample for each replicate. 120ul of Life Technologies Protein A Dynabeads (10002D) were prepared for each ChIP sample by washing beads in 1ml of BSA solution (5%BSA/PBS) at 4°C for 20min. Wash repeated for a total of three times, using a magnet to collect beads and remove BSA solution between washes. After third wash beads were resuspended in 1ml BSA solution, 250ul of which was set aside to preclear the chromatin. 5ul of H2Bub1 antibody (Cell Signaling 5546), and 250ul of fresh BSA solution was added to the remaining 750ul of Dynabeads, and incubated overnight at 4°C. Sheered chromatin samples were pre-cleared by incubating with the 250ul aliquot of Dynabeads resuspended in 2ml of Dilution Buffer (20mM Tris pH=8, 2mM EDTA, 150mM NaCl, 1% Triton X-100, 1:50 Roche cOmplete protease inhibitor) for 1 hour at 4°C. Next the antibody-Dynabead conjugate was rinsed three times in BSA solution, resuspended in 150ul of Dilution Buffer, and added to the pre-cleared chromatin for a 36 hour incubation at 4°C. Beads were then collected on a magnet and washed three times with 1ml of LiCl wash buffer (1% w/v Na-Deoxycholate, 500mM LiCl, 1% NP40 Substitute, 100mM Tris pH=7.5), and twice with TE buffer (10mM Tris-HCl, 1mM EDTA). After removing TE buffer beads were suspended in 75ul of SDS Elution Buffer (1% w/v SDS, 0.1M NaHCO_3_). Samples were then incubated at 37°C for 15 minutes with 1000 RPM mixing. This step was repeated once and the eluates were combined. To reverse crosslinking, input and immunoprecipitated samples were diluted up to 200ul with SDS Elution Buffer, NaCl was added to a final concentration of 200mM, and Proteinase-K to a final concentration of 0.2mg/ml. Samples were then incubated at 65°C overnight. The next day RNase-A was added to a final concentration of 1mg/ml, and incubated at 37°C for 15 minutes. DNA was then purified using a Zymo DNA Clean and Concentrator column according to manufacturer instructions, and eluted in 100ul of H_2_O. NGS sequencing adapter addition and multiplex indexing of DNA samples was achieved using the KAPA Hyper Prep Kit (KAPA Biosystems KK8502) according to manufacturer instructions. 25ng of DNA was used as input. ChIP and input reads were aligned to mm10 using Bowtie2^48^, and reads for each gene were extracted using Bedtools^49^. Reads for each gene, from NCBI RefSeq first transcription start site to last transcription stop site, were summed and normalized to get an RPKM value that served as an H2Bub1 binding score. Input values were subtracted from ChIP values for the final score.

## Supporting information

supp_table1_summary

supp_table2-Screen_results_gRNA

supp_table2-Screen_results_MaGeCK

supp_table3-RNAseq

supp_table4-Custom_gene_sets

supp_table5-GSEA_down

supp_table5_GSEA_up

Supp_table6-gRNA_library

supp_table7-Oligos

supp_table8-Antibodies

## ACKNOWLEDGEMENTS

The authors thank the Dana-Farber Flow Cytometry Core for assistance with cell sorting, and thank Eswar Prasad for guidance related to gRNA reverse transcription. NJV was supported by T32HL007572, F32HL13423501, BCH Kaplan Fellowship, and K99HL143194. The content is solely the responsibility of the authors and does not necessarily represent the official views of the National Heart, Lung, and Blood Institute or the National Institutes of Health. Portions of this research were conducted on the O2 High Performance Compute Cluster, supported by the Research Computing Group, at Harvard Medical School. See http://rc.hms.harvard.edu for more information.

## AUTHOR CONTRIBUTIONS

NJV and WTP conceived and designed the study. NJV executed most experiments. JYL, WG, YZ, and JK generated plasmids, viruses and other necessary reagents, and assisted with processing tissues and cells. YG assisted with experimental design. PZ conducted wildtype CM specific P0 and 4wk RNA-sequencing. QM conducted echocardiography. NJV, IS, SS, GCY, and WTP analyzed the data. NJV and WTP wrote the manuscript.

---

**Supplementary Figure 1.**
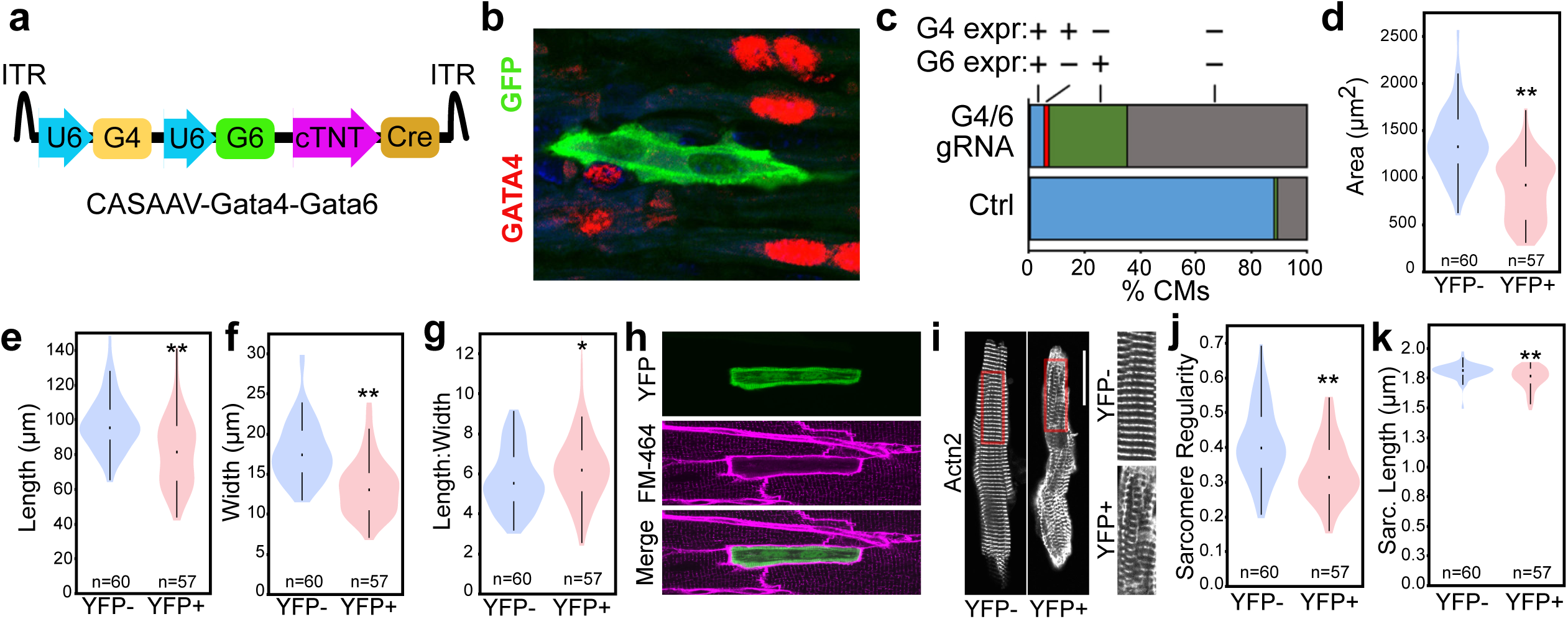
GATA4 and GATA6 are required for normal CM maturation. **a,** CASAAV-Gata4-Gata6 vector. AAV genome encoded CM specific Cre and gRNAs that target Gata4 and Gata6. We administered CASAAV-Gata4-Gata6 at a low dose to postnatal day 1 (P1) mouse pups carrying Cre-activatable Cas9-2A-GFP and *Myh7^YFP^* (R26^Cas9-GFP;^ *Myh7^YFP^*). **b,** GATA4 and GATA6 depletion efficiency. We fixed hearts at P10 for GATA4 and GATA6 immunostaining. Representative section stained for GATA4 is shown. GFP marked transduced cells. **c,** Quantification of GATA4 and GATA6 depletion. Greater than 90% and 60% of the transduced CMs lost GATA4 and GATA6 immunoreactivity, respectfully. **d-g,** Quantification of size and dimensions in isolated P28 CMs, when maturation is normally largely complete. **h,** Imaging of the T-tubule network by FM4-64 staining showed dramatic defects. **i,** Representative image of α-actinin (ACTN2) immunostaining. **j,** Quantification of sarcomere organization. **k,** Quantification of sarcomere spacing.

**Supplementary Fig. 2:**
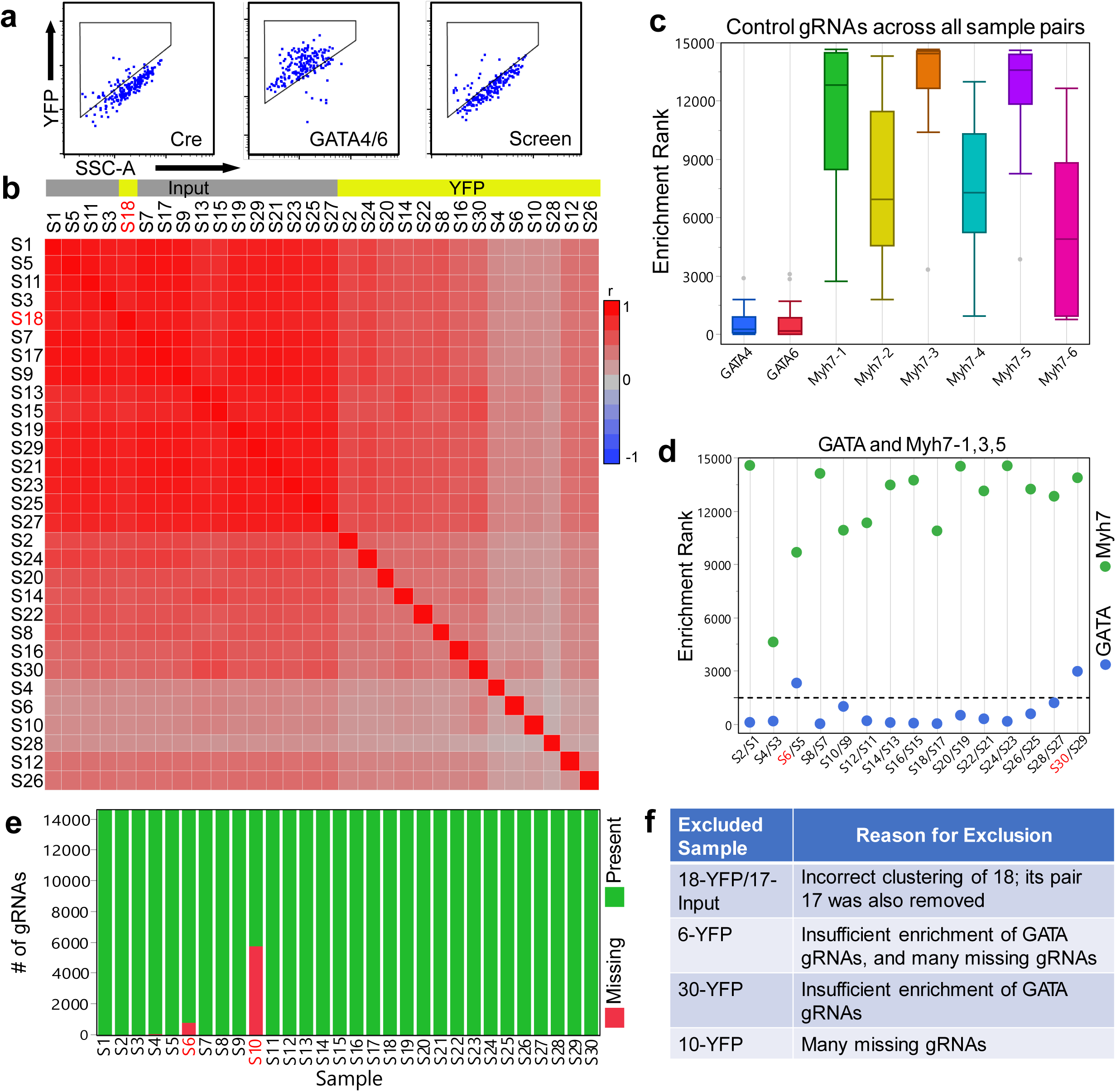
CRISPR screen quality control metrics. **a,** Representative FACS plots showing YFP expression in CMs transduced with Cre, GATA4/6, or gRNA library. **b,** All screen samples, clustered by r correlation. **c,** Enrichment of positive and negative control gRNAs across all samples, where smaller rank indicates higher enrichment. **d,** Average enrichment of GATA gRNAs and efficient Myh7 gRNAs for each sample pair. **e,** The number of gRNAs detected for each sample. **f,** Summary table listing each excluded sample and reason for exclusion. Samples numbers in red font indicates exclusion.

**Supplementary Fig. 3:**
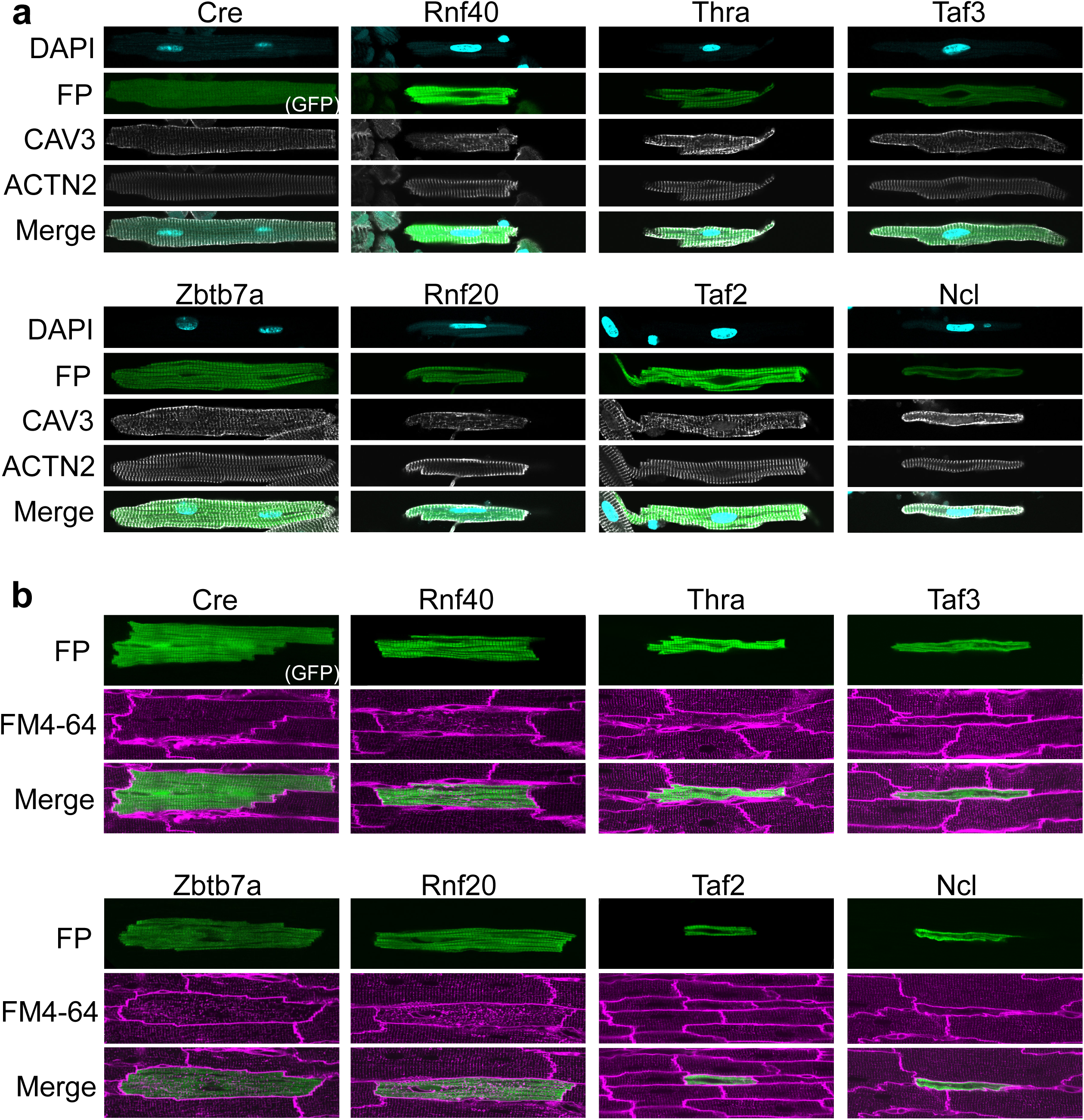
Representative images of CMs featuring depletion of top candidates. **a,** DAPI, Myh7, Caveolin-3 and sarcomeric α-actinin immunostaining of a representative cell for each candidate. **b,** In situ T-tubule imaging of a representative cell for each candidate. FP = fluorescent protein. GFP for Cre, Myh7^YFP^ for candidates.

